# Amyloid precursor protein interacts with the mitochondrial phosphatase PGAM5 and regulates mitochondrial respiration

**DOI:** 10.64898/2026.01.20.700642

**Authors:** Kriti Shukla, Zhi Zhang, Kendra S. Plafker, Satoshi Matsuzaki, Casandra Salinas-Salinas, Yvonne Thomason, Samah Houmam, Dylan Barber, Anna Faakye, Kenneth M. Humphries, Scott M. Plafker, Jialing Lin, Heather C. Rice

## Abstract

Amyloid Precursor Protein (APP) has been reported to partially localize to mitochondria, and mitochondrial dysfunction is a key feature of Alzheimer’s disease; however, the mechanisms linking APP to mitochondrial functions remain incompletely defined. In this study, we identified an interaction between APP and phosphoglycerate mutase family member 5 (PGAM5), a mitochondrial protein phosphatase. We confirmed their endogenous interaction in mouse brain tissue and determined that APP and PGAM5 are both present at mitochondria-ER contact sites (MERCS) and. Using in vitro binding assays, we demonstrate a direct interaction between the linker region of APP and a region of PGAM5 that includes the Kelch-like ECH-associated protein 1 (Keap-1) binding domain. PGAM5 is known to anchor a portion of Nuclear factor erythroid 2 p45-related factor 2 (Nrf2) through Keap1 at the outer mitochondrial membrane and regulates mitochondrial respiration and stress responses. We found that the Nrf2-regulated genes Hmox1 (Heme oxygenase-1) and Nqo1 (NADH:quinone oxidoreductase 1), which are involved in mitochondrial respiration, are downregulated in APP KO astrocytes. Accordingly, mitochondria isolated from the brains of APP knockout (KO) mice have impaired substrate-specific respiration and electron transport chain (ETC) function. Together, these findings suggest that APP supports mitochondrial respiration by binding to PGAM5 and modulating Keap1-Nrf2 signaling.

## Introduction

Mitochondrial dysfunction is a prominent feature of Alzheimer’s disease (AD) and is thought to contribute to disease progression^1–4^. Multiple studies report increased oxidative stress^5,6^, decreased ATP levels^7^, and impaired brain metabolism in AD^8–10^. Amyloid Precursor Protein (APP) is the parent protein of Aβ peptides that aggregate to form plaques in Alzheimer’s disease.^11,12^ Although primarily localized to the secretory pathway, cell surface, and endocytic compartments, a portion of APP has been shown to be present at mitochondria and mitochondria-ER contact sites (MERCS)^13–16^, which are also sites of APP processing^17,18^. Several studies have suggested a role of APP in mitochondrial functions^19–22^. Notably, APP alters mitochondrial mass^23^, bioenergetics^20,24^, mitophagy^25,26^ and mitochondrial respiration^26,27^. The mechanisms by which APP affects mitochondrial function remain unclear, but understanding its protein-protein interactions at the mitochondria can help define its role^28^.

Phosphoglycerate mutase family member 5 (PGAM5) emerged as a candidate interactor of the APP extracellular domain in an unbiased proteomics screen performed by Rice et al.^28^. PGAM5 is a mitochondrial phosphatase, with substrates that include B-cell lymphoma-extra large (Bcl-XL)^29^, Dynamin-related protein 1 (DRP1)^30,31^, FUN14 domain-containing protein 1 (FUNDC1)^32^ and Mitofusin 2 (MFN2)^33^, which are involved in mitochondrial mobility, mitophagy, redox homeostasis, respiration, or cell death.^29–32,34–36^. PGAM5 also regulates mitochondrial respiration and stress response through the Keap1-Nrf2 pathway^36–41^. Under homeostatic conditions, Nrf2 is constitutively translated, ubiquitylated by Keap1/Cul3, and degraded by the proteasome^42,43^. In response to oxidative stress, modification of cysteines on Keap1 render it enzymatically inactive leading to a halt of the constitutive degradation of Nrf2. Nrf2 accumulates in the cytoplasm and then translocates to the nucleus, where it induces the transcription of antioxidant genes (e.g. *Hmox1 and Nqo1*) to restore redox homeostasis and support mitochondrial respiration^36,42,43^. A subset of Nrf2 is tethered to the outer mitochondrial membrane through a ternary complex with PGAM5 and Keap1. Knockdown of PGAM5 or Keap1 activates Nrf2 target gene expression ^36,40^. The PGAM5-Keap1-Nrf2 complex has been implicated in both stress-induced mitochondrial retrograde trafficking^37^ and mitochondrial respiration^36^.

In this study, we establish an interaction between APP and PGAM5 that is mediated by a region of PGAM5 that includes the Keap1-binding domain. Consistent with disrupted PGAM5-Keap1-Nrf2 signaling, expression of the Nrf2 target genes *Hmox1 and Nqo1* are reduced in APP KO astrocytes. We show that mitochondria from APP KO mouse brains have impaired substrate-specific respiration and electron transport chain (ETC) function. Together, these findings suggest a role for APP in regulating mitochondrial respiration by binding to PGAM5 and regulating Keap1-Nrf2 signaling pathway.

## Results

### APP and PGAM5 interact endogenously and are both present at mitochondria-ER contact sites

To investigate a potential mechanism underlying APP function in mitochondria, we followed up on a prior proteomics screen performed by Rice et. al. ^28^ identifying the mitochondrial phosphatase PGAM5 as a candidate interactor of the APP ectodomain^4^. To determine whether APP and PGAM5 interact endogenously, PGAM5 was immunoprecipitated (IPed) from brain extracts of wild-type (WT) mice or PGAM5 knockout (KO) mice, as a negative control. APP co-IPed with PGAM5 in WT but was not detected in IPs from PGAM5 KO extracts, indicating that APP and PGAM5 interact endogenously in mouse brain (Figure 1A). To confirm that a portion of APP and PGAM5 localize to common subcellular compartments, we performed subcellular fractionation of WT mouse brains to isolate mitochondria, ER, and mitochondria-ER contact sites (MERCS). Both young (3mo) and old (12mo) mice showed similar subcellular distribution of APP and PGAM5, so both ages were plotted together. Consistent with the literature, APP was found to localize at the ER but interestingly, a significant amount was also present at MERCS (Figure 1B,C). On the other hand, PGAM5 was also found to be present at MERCS while predominantly being present at the mitochondria (Figure 1B,D). This data showed that both APP and PGAM5 can be found at MERCS (Figure 1B-D), suggesting MERCS as a potential site for their interaction.

**Figure 1.**
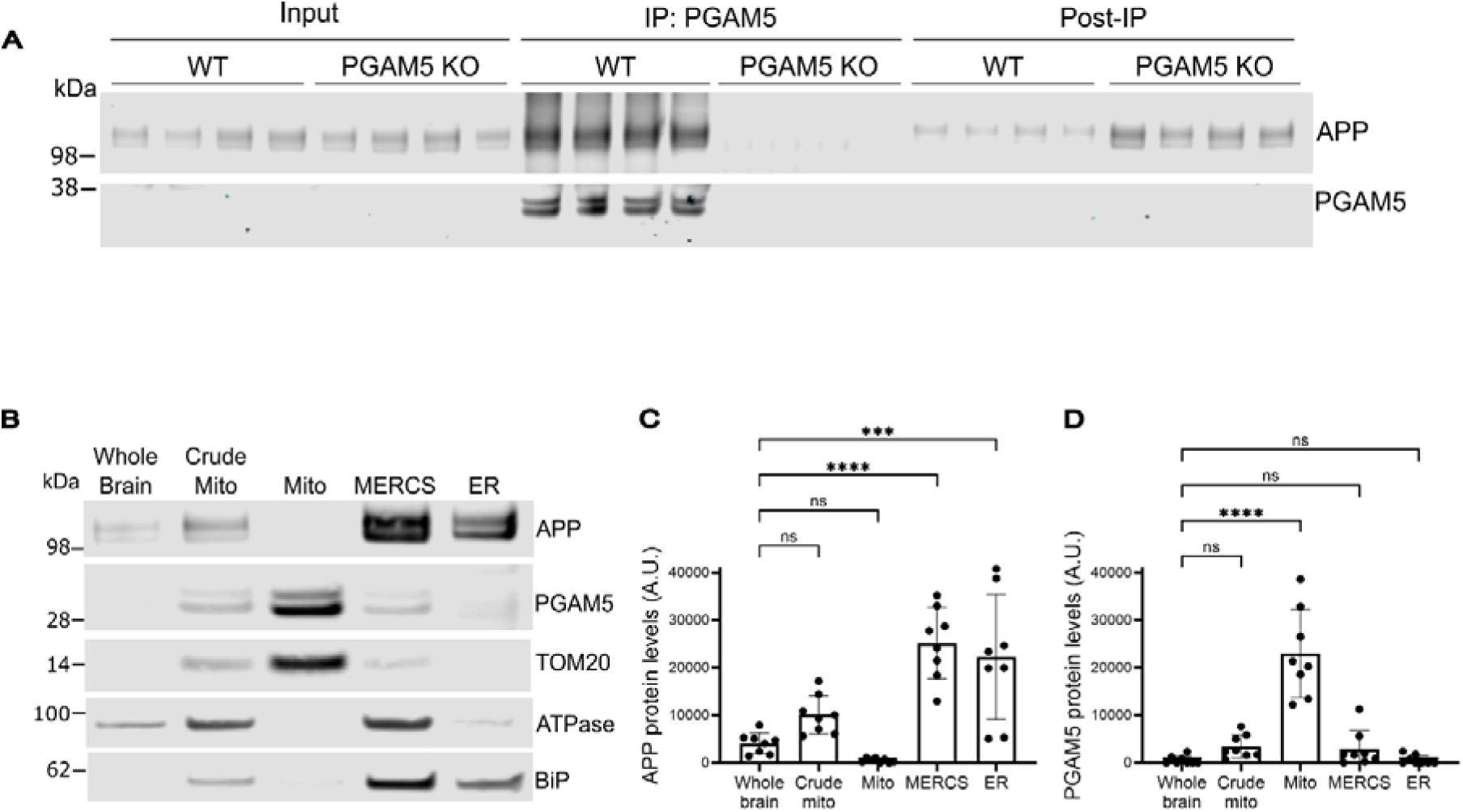
APP and PGAM5 interact endogenously and are both present at mitochondria-ER contact sites. (**A**). Western blot showing immunoprecipitated PGAM5 and co-immunoprecipitated APP in WT and PGAM5 KO (4mo, males) mouse brains. **(B).** Western blot showing subcellular localization of PGAM5 and APP in WT (C57Bl6, 3-12 months old, females) mouse brains. The cellular compartments shown are Whole brain, Crude mitochondria, Mitochondria, Mitochondria-ER contact sites (MERCS) and Endoplasmic reticulum (ER). The purity of each sample was validated by its marker protein (TOM20 (Translocase of Outer Mitochondrial Membrane 20), Mitochondria; ATPase (Adenosine Triphosphatase), MERCS; BiP (Binding Immunoglobulin Protein), ER). **(C-D).** Graphs showing western blot quantification indicating levels of C. APP protein and D. PGAM5 protein. Values shown are band intensity means ± SD (*n* =7). One-way ANOVA tests were performed for each graph, *** (*P* < 0.01), **** (*P* < 0.001), ns (non-significant).

### APP interacts with a region of PGAM5 containing the Keap1 binding domain

Next, we used *in vitro* binding assays to determine whether APP and PGAM5 can directly interact and to narrow the binding regions responsible for this interaction. Since both APP and PGAM5 full-length proteins have transmembrane domains (TM), we used the recombinant soluble APP (sAPPα, aa1-496) (Figure 2A) and truncated PGAM5 constructs (Figure 2B). Amino acids 1-54 (TM domain) of PGAM5 were removed to make PGAM5Δ54 while PGAM5Δ90 had additional 36 amino acids (containing the multimerization motif;MM and Keap1-binding domain;KBD) removed^4734,36,48^. We used Protein G Sepharose beads to pull down Fc-tagged sAPPα incubated with either PGAM5Δ54 or PGAM5Δ90. PGAM5Δ54 but not PGAM5Δ90 was pulled-down with sAPPα-Fc, indicating that the interaction requires the N-terminal region of PGAM5 containing the KBD and MM (Figure 2C). Isothermal titration calorimetry (ITC) with sAPPα (without an Fc tag) and PGAM5Δ54, revealed a dissociation constant (K_d_) of 3.45 µM and a binding stoichiometry of N = 0.114 (Figure 2D). Then, to narrow down the domain in APP required for binding to PGAM5, we used Protein G Sepharose beads to pull down individual Fc-tagged domains of APP (Growth factor-like domain;GFLD-Fc, Copper binding domain;CuBD-Fc, Extension domain;ExD-Fc, Acidic domain;AcD-Fc and Extracellular 2;E2-Fc) incubated with PGAM5Δ54. PGAM5Δ54 was most efficiently pulled-down with ExD-Fc followed by AcD-Fc (Figure 2E-F). Together, these results suggest that the linker region of APP (constituted by AcD and ExD) and the N-terminal amino acids (aa55-89) of PGAM5 (containing Keap1 binding domain) are critical for binding between these two proteins.

**Figure 2.**
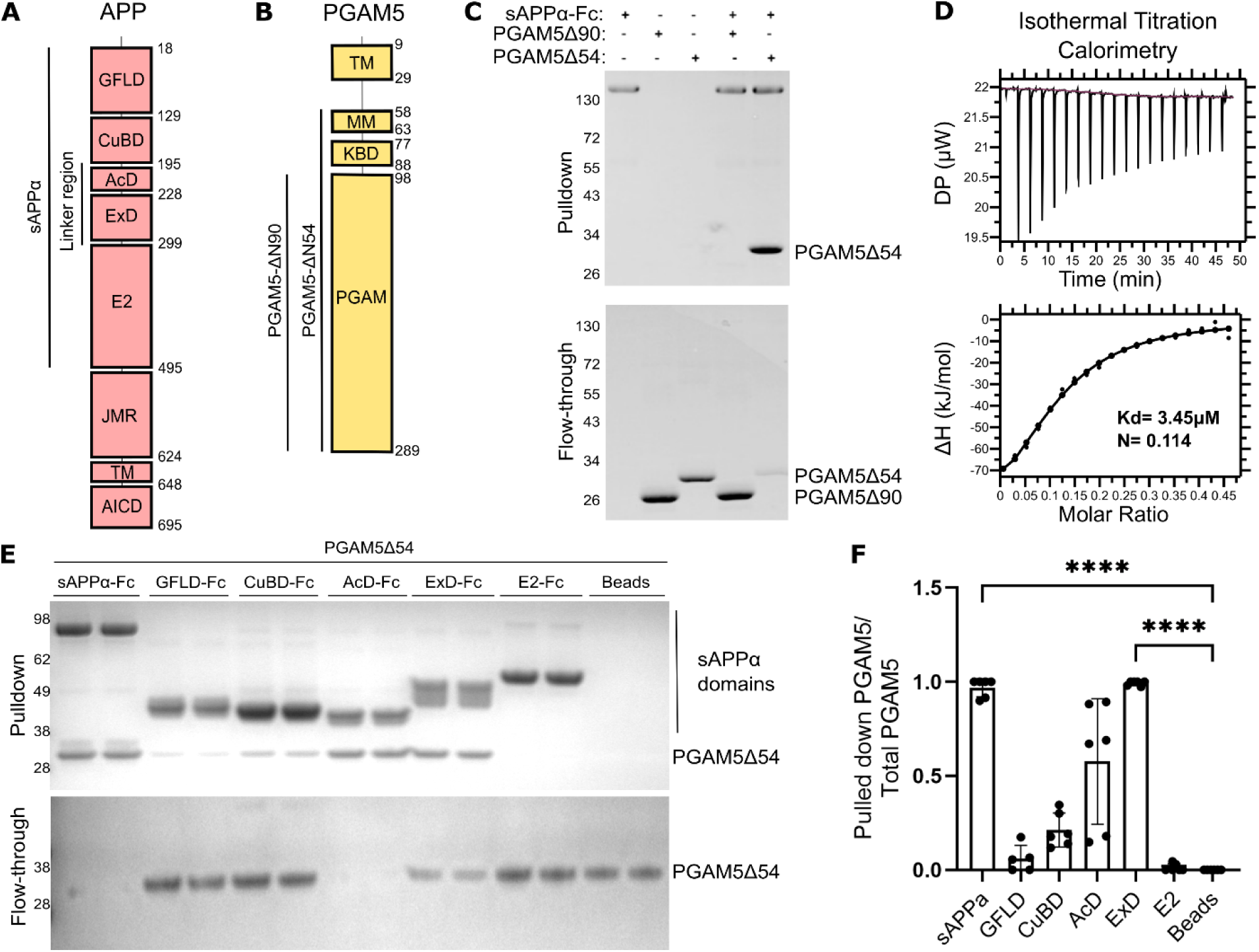
APP interacts with a region of PGAM5 containing the Keap1 binding domain. (**A-B**). Schematics showing domains (GFLD; Growth-like factor domain, CuBD; Copper-binding domain, AcD; Acidic domain, ExD; Extension domain, E2, JMR; Juxtamembrane Region, TM; Transmembrane, AICD; APP intracellular domain, MM; Multimerization motif, KBD; Keap1 binding domain, PGAM; Phosphoglycerate mutase) and recombinant protein constructs (sAPPα, PGAM5-Δ90 and PGAM5-Δ54) of APP (pink) and PGAM5 (yellow). **B.** Coomassie stained SDS-PAGE gels of bound and unbound material from Fc-pulldowns of sAPP-Fc with PGAM5Δ54 or PGAM5Δ90 purified proteins, **C.** Binding curve from ITC done using the sAPPα and PGAM5-Δ54 purified proteins (n=3). **(D-E).** D. Coomassie stained SDS-PAGE gels of bound and unbound material from Fc-pulldowns of sAPP-Fc, GFLD-Fc, CuBD-Fc, ExD-Fc, AcD-Fc, E2-Fc and beads with PGAM5Δ54 purified protein and E. Quantification of pull-down assays showing net PGAM5Δ54 bound and pulled down with each Fc-tagged sAPPα domain. Values shown are means ± SD (*n* =5 – 6) and the statistics were performed using Dunnett’s multiple comparison test. ***** (*P* < 0.0001).

### Transcription of Nrf2-regulated genes is altered in APP KO astrocytes

To understand the functional implications of binding of APP to the N-terminal region of PGAM5 containing the Keap-1 binding domain, we investigated whether the Keap1-Nrf2 pathway is affected by the presence of APP. We examined Nrf2-dependent gene expression in primary astrocytes isolated from the brains of WT or APP KO mice. Astrocytes were chosen based on prior literature demonstrating a critical role for APP in mitochondrial function in astrocytes^44^. Total RNA was isolated from WT or APP KO astrocytes and transcript levels of established Nrf2 target genes (*Hmox1, Nqo1, Gclc, Gclm, and Txnrd1*) were quantified by qPCR. Both *Hmox1* and *Nqo1* transcripts were significantly reduced in APP KO astrocytes compared to WT controls. No significant differences were observed for *Gclc*, *Gclm*, or *Txnrd1* (Figure 3).

**Figure 3.**
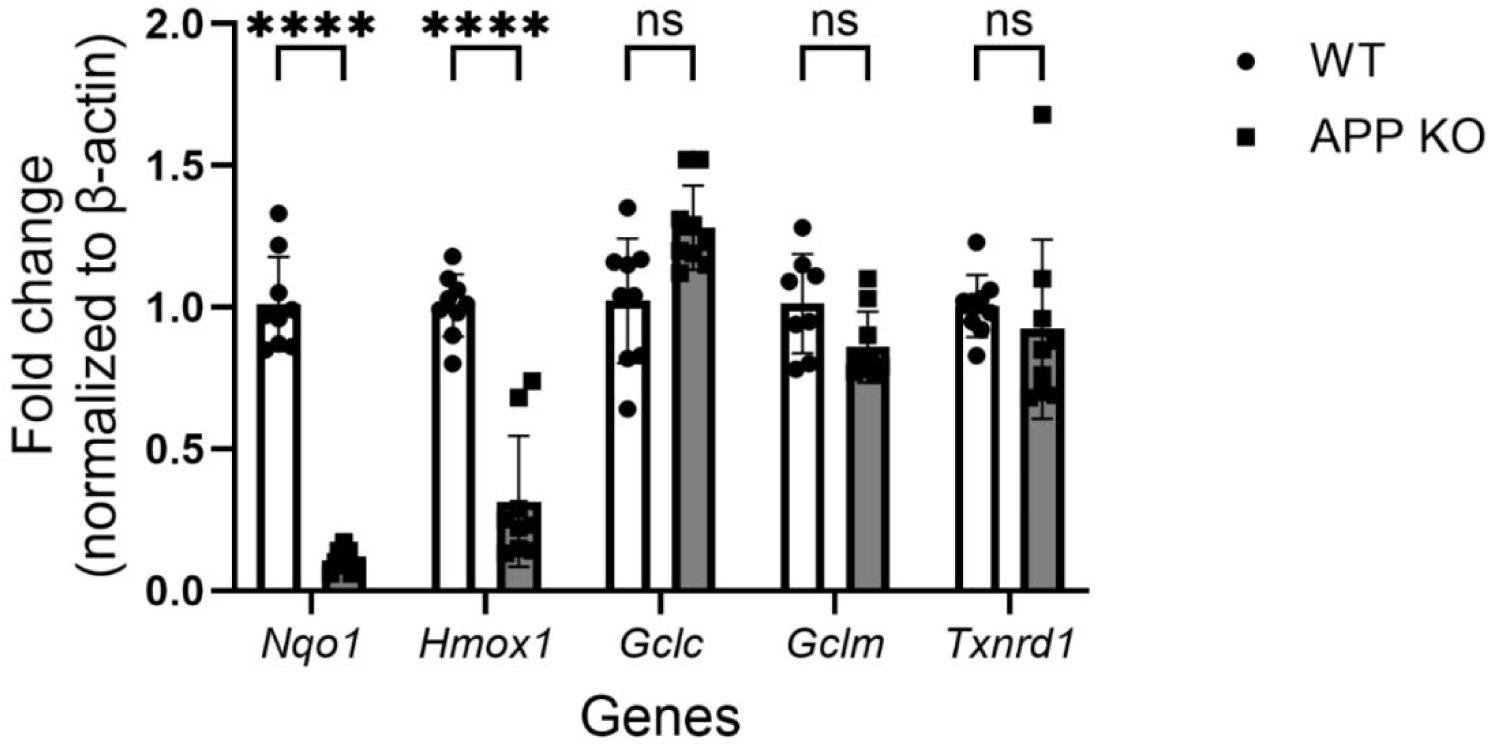
APP affects Nrf2-regulated gene expression. Plot showing relative mRNA expression of Nrf2-regulated genes in cultured primary astrocytes isolated from WT and APP KO mouse brains, measured by RT-qPCR. Values shown are the means ± SD (*n* = 8-9) and the statistics were performed using two-way ANOVA, followed by Tukey’s multiple comparison test. ***** (*P* < 0.0001), ns (non-significant).

### Mitochondria isolated from APP KO mouse brains have impaired respiration

Since the basal expression of *Nqo1* and *Hmox1* was decreased in APP KO conditions and both these genes are essential for maintaining redox homeostasis that permit efficient NADH oxidation at Complex I^45,46^, we investigated if loss of APP also affects mitochondrial respiration and ETC. We performed respiratory measurements in freshly isolated mitochondria from WT vs APP KO mouse brains. APP KO mitochondria showed significantly decreased pyruvate (Figure 4A) and glutamate-mediated (Figure 4B) respiration with no changes observed in succinate-mediated respiration (Figure 4C) as compared to WT, indicating an impairment in maximal NADH-linked mitochondrial respiration whereas Complex II–dependent respiration (succinate) remains intact. Additionally, we measured the effect of APP KO on Complex I activity, the rate limiting step of the ETC and NADH oxidase activity as a measure of total ETC activity. Our findings showed a significant decrease in NADH oxidase activity in APP KO mitochondria (Figure 4D) and a decreasing but non-statistically significant trend in complex I activity (Figure 4E). Taken together, these findings highlight a crucial physiological role of APP in maintaining Complex I function and NADH-linked respiration in brain mitochondria.

**Figure 4.**
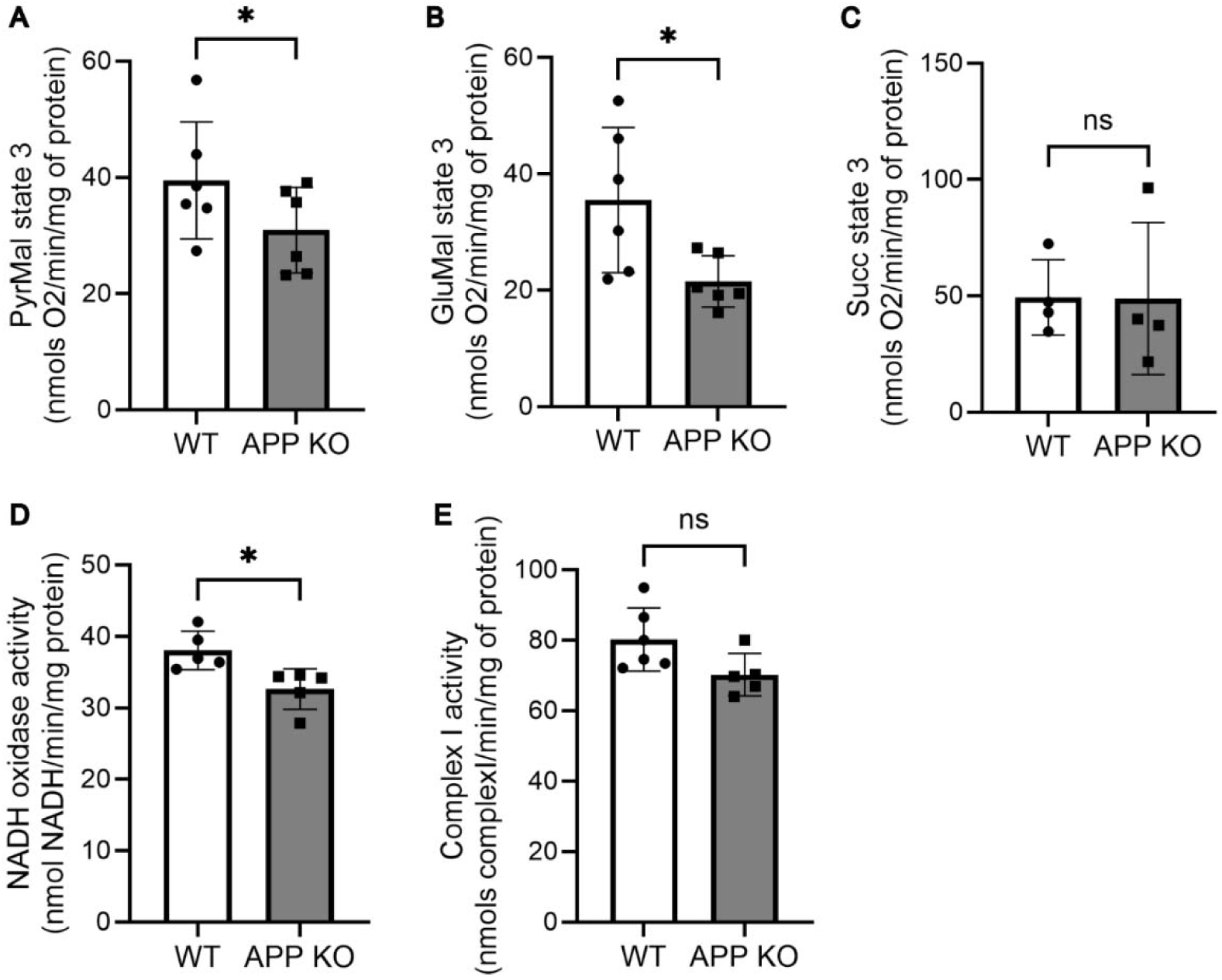
Isolated mitochondria from APP KO mice brain have impaired respiration. (**A-C**). Graphs showing maximum respiration by utilizing different oxidizable substrates (A. Pyruvate/malate, B. glutamate/malate, C. Succinate), in the mitochondria isolated from 6-8 months old APP KO mouse brains. Values shown are the means ± SD (*n* =4 – 6). Two-tailed paired t-tests were performed for each graph. **(D-E).** Graphs showing impaired ETC in APP KO mitochondria as indicated by D. decreased Complex I activity and E. NADH oxidase activity. Values shown are means ± SD (*n* =5 – 6). Two-tailed paired t-test was performed for each graph, * (*P* < 0.05), ns (non-significant).

## Discussion

In this study, we identify the mitochondrial protein phosphatase PGAM5 as an interactor of APP. Both APP and PGAM5 were found to interact endogenously and both were present at MERCS. Interestingly, the amount of APP present at MERCS was comparable to the amount at ER, suggesting a crucial role of APP at these inter-organelle sites. Previous studies have identified MERCS as an APP processing site ^17,18^, and our findings suggest a functional role for APP localized to MERCS^23,24^. Our ITC data showed binding between APP and PGAM5 with a binding stoichiometry (N) of 0.114, which is consistent with a dodecameric PGAM5 complex associated with an APP dimer. APP is known to form homodimers through its E1 and transmembrane domains, which has been implicated in its processing and function^47^. PGAM5, on the other hand, assembles into higher-order oligomers, predominantly forming dodecamers as well as dimers, that are critical for its functions^30,32,48^ The APP-PGAM5 interaction requires a region of PGAM5 containing the Keap1-binding domain, suggesting that APP may compete with Keap1 for its binding to PGAM5. PGAM5 is known to tether Nrf2 at the outer mitochondrial membrane through Keap1 and regulates the amount of Nrf2 translocating into the nucleus for gene expression^40^. Consistent with this, we find that transcription of Nrf2 target genes, *Nqo1* and *Hmox1,* is significantly reduced in APP KO primary astrocytes under physiological conditions. Both NQO1 and HMOX1 are critical components of the Nrf2 antioxidant network that preserve mitochondrial redox homeostasis and regulate respiration^49–51^. NQO1 limits quinone redox cycling and supports respiration by maintaining NAD⁺/NADH balance^52,53^, while HMOX1 helps mitigate oxidative stress and supports ETC^54,55^. Taken together, these findings suggest that the APP-PGAM5 interaction at MERCS may regulate mitochondrial respiration and stress response through modulation of Keap1-Nrf2 signaling.

We further show that APP deficiency leads to a selective reduction in glutamate– and pyruvate-driven respiration. This substrate-specific defect points to a dysfunction upstream of or at Complex I, rather than a global failure of the electron transport chain or mitochondrial integrity. Consistent with this interpretation, we observe a reduction in NADH oxidase activity and a downward trend in Complex I enzymatic activity in APP KO mitochondria. Together, these results position APP as a regulator of NADH-linked respiration in the brain and suggest that impaired NADH-dependent respiration may not only limit electron transport capacity but also promote oxidative imbalance and bioenergetic insufficiency. In this context, diminished Complex I function could enhance electron leak and reactive oxygen species generation while simultaneously constraining ATP synthesis. When considered alongside previous evidence that manipulation of APP alters ETC activity, ATP levels, and mitochondrial biogenesis^26,56^, our findings strengthen the conclusion that APP is an essential contributor to mitochondrial bioenergetics.

Based on our findings, we propose a model in which APP helps maintain Nrf2-dependant gene regulation by competing with Keap1 to bind to PGAM5 at MERCS. In WT conditions, APP binding to PGAM5 may promote displacement of Keap1-bound Nrf2, enabling Nrf2 accumulation at the cytoplasm and eventual nuclear translocation that in turn leads to gene expression (Figure 5A). On the other hand, in the absence of APP, Keap1-bound Nrf2 is constantly tethered to PGAM5 and is unable to translocate to the nucleus to turn on the gene transcription and hence the expression of genes required to maintain mitochondrial respiration and redox balance is reduced (Figure 5B). This model provides a mechanistic basis for mitochondrial bioenergetic deficits often observed upon APP loss^61–63^ and warrants future studies to determine whether mitochondrial dysfunction observed in AD arises from loss of normal APP function.

**Figure 5.**
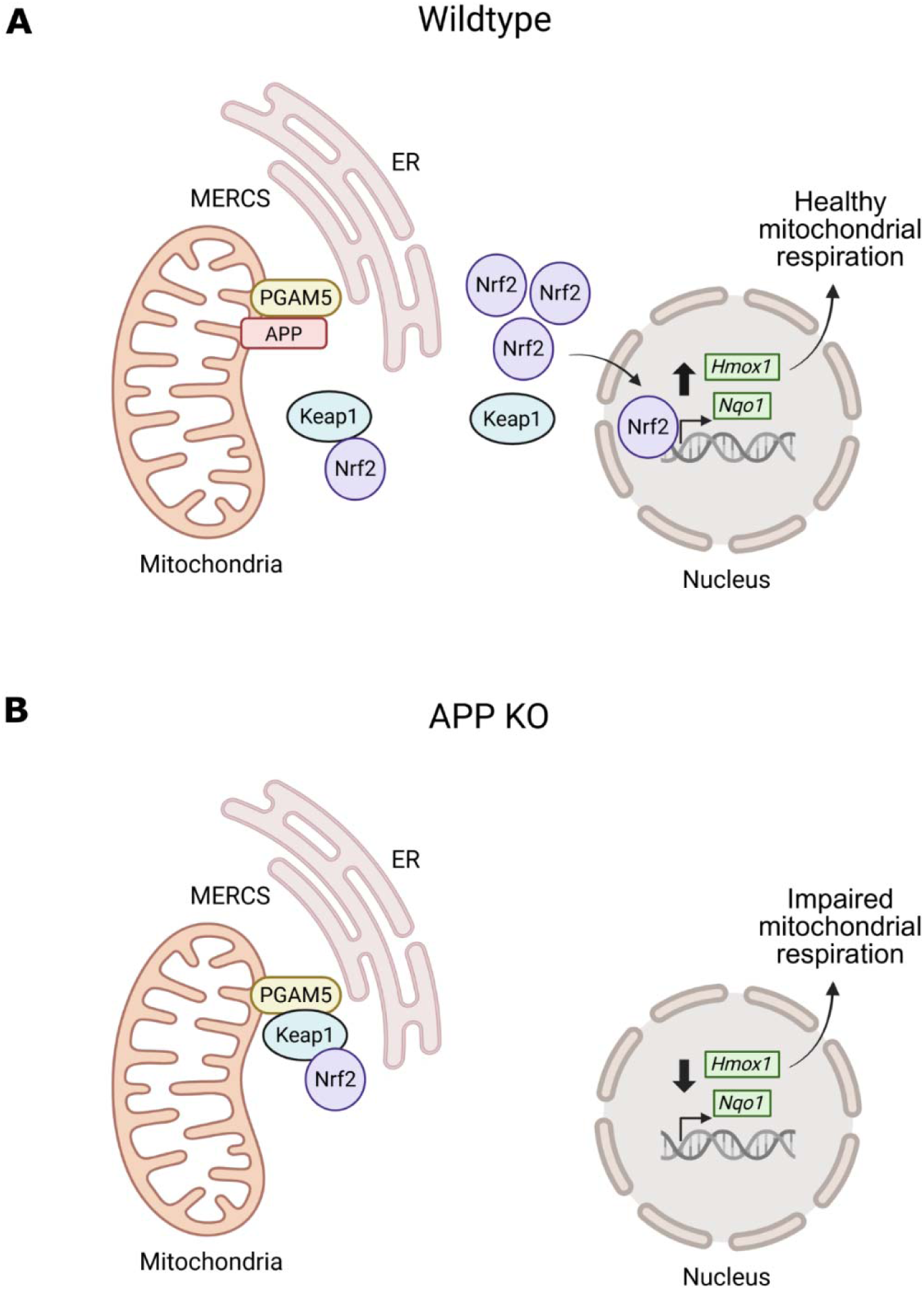
Proposed model by which APP affects mitochondrial respiration. **A.** In WT cells, APP localized to MERCS can bind to PGAM5 and displace Keap1. This releases Keap1-bound Nrf2 to the cytoplasm where Nrf2 dissociates from Keap1, accumulates, and subsequently translocates to the nucleus to activate transcription of genes that maintain normal mitochondrial respiration. **B.** In APP KO cells, Nrf2 remains bound to PGAM5 at the mitochondria, leading to reduced Nrf2 translocation to the nucleus and reduced transcription of its target genes. This reduced Nrf2 signaling in the absence of APP leads to impaired mitochondrial respiration. The schematic was created in BioRender. Rice, H. (2026) https://BioRender.com/ftsycbt.

## Experimental procedures

(see Supporting information for details)

## Mouse models

All experiments were conducted following a protocol approved by the Oklahoma Medical Research Foundation’s (OMRF) Institutional Animal Care and Use Committee prior to any animal work.

## Co-immunoprecipitation, Western blotting, Protein pull-down assays

see Supporting information for details

## Subcellular fractionation

Mouse brain mitochondria and mitochondria-ER contact sites (MERCS) were isolated as previously described ^57^.

## Plasmids

APP plasmids were provided by Dr. Bart De Strooper and Dr. Joris de Wit (VIB-KU Leuven, Belgium)^28^. PGAM5 plasmids were provided by Dr. Apirat Chaikuad (Institute for Pharmaceutical Chemistry, Johann Wolfgang Goethe-University and Buchmann Institute for Molecular Life Sciences, Germany)^48^.

## Purification of recombinant soluble APP proteins

APP fragments with a C-terminal Fc tag were expressed by transfection in HEK293T cells and purified using Protein-G Sepharose 4 Fast Flow resin.

## Purification of recombinant soluble PGAM5 proteins

PGAM5 plasmids were expressed in BL21 Rosetta competent *E. coli* cells using 0.5mM IPTG. The proteins were purified using Nickel(II)-NTA agarose beads.

## Isothermal Titration Calorimetry

All ITC experiments were performed on a MicroCal PEAQ-ITC system at OU Protein Production and Characterization Core. The raw ITC data were fitted to a single binding site model using the MicroCal PEAQ-ITC analysis software provided by the manufacturer.

## Primary Mouse Astrocyte Culture

Primary astrocytes were isolated from whole brains of postnatal day 0–5 (P0–P5) C57BL/6J or APP KO mice using a differential adhesion–based protocol^58^.

## qPCR

Total RNA extracted from primary mouse astrocytes was used to synthesize cDNA. Real-time quantitative PCR was performed using PowerUp™ SYBR^®^ Green PCR Master Mix (Applied Biosystems) and the primers for the genes– *Hmox1, Nqo1, Gclc, Gclm*, *Txnrd1* and *Actb.* The mRNA levels of target genes were normalized to *ActB* by untreated WT sample values.

## Respiratory measurements with isolated mitochondria

Mouse brain mitochondria were isolated and mitochondrial respiration measurements were performed according to established protocols.^57,59^

## ETC enzyme activity assays

NADH oxidase activity was measured as previously described.^60^

## Data Availability

All data described in the manuscript are contained within the manuscript.

## Supporting Information

This article contains supporting information^28,48,57–61^

## Acknowledgements

This work was supported by the National Institutes of Health (R35GM142726, P20GM125528 to H.C.R.), Institutional Development Award (IDeA) from the National Institute of General Medical Sciences (NIGMS) of the National Institutes of Health (NIH) (P20GM103640 to J.L. as a junior investigator), and the Presbyterian Health Foundation (Pilot Research Funding to H.C.R.; Bridge Funding to J.L..). APP plasmids were provided by Dr. Bart De Strooper and Dr. Joris de Wit (VIB-KU Leuven, Belgium)^28^. PGAM5 plasmids were provided by Dr. Apirat Chaikuad (Institute for Pharmaceutical Chemistry, Johann Wolfgang Goethe-University and Buchmann Institute for Molecular Life Sciences, Germany)^48^. The ITC experiments were done at OU Protein Production and Characterization Core facility, supported by IDeA from the NIGMS of NIH (P20GM103640 and P30GM145423), the OU Vice President for Research and Partnerships, and the OU College of Arts and Sciences. PGAM5 KO mouse strain was provided by Dr. Yishi Jin with permission from Dr. Michael Lenardo who originally developed the mice^61^. Data analysis was supported by the OMRF Center for Biomedical Data Sciences. Figure 5 schematic was created in Biorender-https://BioRender.com/ftsycbt

