## Supplemental Methods for "Amyloid precursor protein interacts with the mitochondrial phosphatase PGAM5 and regulates mitochondrial respiration"

**Materials include:**

Extended experimental procedures

**Mouse models**

Wild-type mouse strain used in this study was C57/Bl6 (starin 000664, The Jackson Laboratory) and genetically modified mouse strains were APP KO (strain 004113, B6.129S7-App/J, The Jackson Laboratory) and PGAM5 KO (MGI:4432458, Pgam5^tm1a(EUCOMM)Wtsi^ , received by Dr. Yishi Jin, University of California San Diego and originally developed by Dr. Michael Lenardo, National Institute of Health)^61^. All the mice were group-housed in the AALAS-accredited OMRF vivarium on a 14:10 h light/dark cycle with *ad libitum* access to water and food. The homozygosity of the APP KO and PGAM5 KO strains was confirmed by genotyping ear-punch biopsies via Transnetyx. Genotyping of all progenies was performed by Transnetyx Inc., which utilizes duplicate sample processing using real-time PCR, ensuring the accuracy of each mutation. Additionally, Transnetyx stores all samples for a minimum of 4 months, should re-probing be necessary. This ensures that in all studies, authentication of mouse lines is performed before use and at regular intervals by a reputable, outside party. For APP KO model, two TaqMan assays, Neomycin (binds within generic neomycin and detects the mutant allele) and App-3 WT (binds within the region deleted by neomycin and provides zygosity), are used to detect the mutant and WT alleles, respectively. For PGAM5 KO model, the probes Pgam5-1 WT and L1L2-Bact-P TA were used. Each assay runs alongside an internal housekeeping gene, Cjun, to ensure there is enough DNA for a reaction and that all other quality controls are passed.

**Co-immunoprecipitation**

Whole mouse brains were homogenized in T-PER buffer supplemented with protease and phosphatase inhibitors, or previously prepared and aliquoted brain lysates (WT and PGAM5 KO) were thawed on ice. Lysates were pre-cleared with Protein A Sepharose 4 Fast Flow beads (cat. No. P9424, Sigma-Aldrich) (washed 3X in T-PER with inhibitors) for 3 h at 4°C on a nutator. Samples were centrifuged at 3,000xg for 30 s at 4°C, and the supernatant (pre-cleared lysate) was collected. A part of this pre-cleared lysate was saved as input. Rest was incubated overnight at 4°C with 1.5 µg PGAM5 rabbit polyclonal antibody (Abcam, ab126534) and 30 µL washed Protein A Sepharose 4B beads on a nutator. The following day, beads were pelleted (3,000xg, 30 s, 4°C), and the supernatant was collected as the post-IP fraction. The protein present in post-IP sample was acetone precipitated as described below. Beads were washed three times with 200–300 µL T-PER (with inhibitors), rotating 30 min at 4°C for each wash. After the final wash, residual buffer was removed, and beads were resuspended in 4X loading dye supplemented with 10X reducing agent, heated at 95°C for 10 min, and centrifuged to pellet beads.The supernatant was collected as the IP sample. Input and post-IP samples were prepared in parallel and analyzed by Western blot (described below).

**Acetone Precipitation**

Required volume of acetone was placed at -20°C. 4 volumes of this chilled acetone were added to 1 volume of proteins sample in acetone-compatible tube. The tube was vortexed and incubated for 60 minutes at -20°C. Afterwards, the tube was centrifuged for 10 min at 13,000-15,000×g. Supernatant was properly and carefully discarded, being careful not to dislodge the protein pellet. Acetone was allowed to evaporate from the uncapped tube at room temperature for 30 minutes. Making sure not to over-dry the pellet, the pellet was resuspended in the T-Per buffer and 4x dye and 10x reducing agent was added to prepare the samples for Western blotting.

**Western blotting**

Samples obtained from subcellular fractionation were quantified using the BCA protein assay (Thermo Fisher Scientific). 12 µg of protein sample was loaded on each lane, mixed with 10x reducing agent and 4X Laemmli sample buffer, boiled for 5 minutes at 95°C, and resolved on NuPAGE^TM^ 4-12% Bis-Tris Midi gels with MES SDS running buffer. Proteins were transferred onto nitrocellulose membranes (BioRad) using Criterion Blotter (BioRad) at 400mA for 80 minutes. Membranes were blocked with 5% BSA in water for 1 hour at room temperature and then incubated overnight at 4°C with primary antibodies diluted 1:1000 in 0.05% Tween-20 in PBS (PBST)- APPY188 (rabbit, abcam, ab32136), PGAM5 (rabbit, abcam, ab126534), TOM20 (rabbit, Thermo-Fisher, 11802-1-AP), ATPase (mouse, Sigma-Aldrich, 05-369-25UG), BiP (rabbit, abcam, ab108613). After washing three times with PBST, membranes were incubated with secondary antibodies (Alexa Flour^TM^ Donkey anti-Rabbit 800 and Donkey anti-Mouse 680) for 1 hour at room temperature. Protein bands were visualized using Odyssey CLx detection system (Licor). The gel image was converted to grayscale, and the protein and background band intensities were obtained using ImageJ software The background intensity was then used to normalize the protein band intensity, and these values were plotted as protein levels in each subcellular compartment.

**Protein pull-down assays**

3 µg of Fc-tagged sAPPα (~95KD) was incubated with either 3 µg PGAM5-ΔN90 (~25kDa) or PGAM5-ΔN54 (~30kDa) in 500 µL binding buffer (10mM HEPES pH 7.4, 150mM NaCl, 2mM CaCl_2_, 1mM MgCl_2_, 0.1% Tween-20) at 4 °C for 1 hour with gentle rotation. For pull-down, 20ml of protein G Sepharose 4 Fast Flow beads (cat. 17061805, Cytiva), pre-equilibrated with cold binding buffer, were added and incubated for an additional 2 hours at 4 °C. Afterwards, flow-through was collected and the beads were washed 3–4 times with 500µl cold binding buffer followed by centrifugation at 3000rpm for 2 min to remove non-specifically bound proteins. Proteins bound to the beads were eluted in 25ml of SDS sample buffer, supplemented by 10% β-mercaptoethanol, heated at 95[°C for 5 minutes and](https://www.degreesymbol.net/) resolved on 13% SDS-PAGE gels, along with the flow-through samples. Gels were then incubated for 1 hour at room temperature with the Coomassie stain solution followed by destain solution and then washed with deionized water (3 times, 15 min each). The gels were then imaged using white light in Syngene apparatus to visualize the protein bands. Same protocol was followed to narrow down the interacting domain in APP by using 3 µg of Fc-tagged Growth factor-like domain;GFLD, Copper binding domain;CuBD, Extension domain;ExD, Acidic domain;AcD and Extracellular 2;E2, with 3 µg of PGAM5-ΔN54.

**Subcellular fractionation**

Brain mitochondria and mitochondria-ER contact sites (MERCS) were isolated from 3-12 mo old female C57/Bl6mouse brains using a modified version of the protocol by Wieckowski et al. (2009)^57^. Briefly, freshly dissected brains were homogenized in ice-cold isolation buffer containing 0.25 M sucrose, 10 mM HEPES (pH 7.4), and protease inhibitors. For each replicate, the homogenate from 3-5 mouse brains was pooled and was subjected to sequential centrifugation to remove nuclei and debris, followed by differential centrifugation steps to separate crude mitochondria. A Percoll density gradient was used to purify mitochondria, and the post-mitochondrial supernatant was further processed to isolate MERCS. All steps were performed at 4°C to preserve organelle integrity.

**Plasmids**

These were provided by Dr. Bart De Strooper and Dr. Joris de Wit (VIB-KU Leuven, Belgium)^28^. These were previously generated by PCR-amplifying the following regions of mouse APP_695_: sAPPα (aa18-612); GFLD (aa18-128); CuBD (aa129-194); AcD-Exd (aa195-298); ExD (aa195-227); AcD (aa228-298); and E2 (aa299-494). Each of the PCR fragments were inserted in frame between the prolactin signal peptide and human Fc sequence in the pCMV6-XL4 vector using Gibson Assembly (NEB). PGAM5 plasmids were provided by Dr. Apirat Chaikuad (Institute for Pharmaceutical Chemistry, Johann Wolfgang Goethe-University and Buchmann Institute for Molecular Life Sciences,Germany)^48^.

**Purification of recombinant soluble APP proteins**

APP fragments with a C-terminal Fc tag were expressed by transfection in HEK293T cells using polyethylenimine (PEI) (Kyfora Bio 23966) and collected in serum-free Opti-MEM media (Gibco 31985088). For Fc-tagged proteins used for *in vitro* binding assays, the conditioned medium was applied to an affinity column packed with Protein-G Sepharose Fast Flow resin (Cytiva 17061802) at 4 °C at a flow rate ~50mL/hr, washed with 250 mL wash buffer (50 mM HEPES pH 7.4, 300 mM NaCl) and eluted with 10 mL of IgG elution buffer (Pierce). For non-Fc-tagged proteins used in ITC, the conditioned medium was passed through a Protein-G Sepharose Fast Flow column, followed by washing with 250 mL of wash buffer (50 mM Tris pH 8.0, 450 mM NaCl, 1 mM EDTA). The Fc tag was cleaved by overnight incubation with GST-tagged 3C PreScission Protease (Cytiva 27084301) in cleavage buffer (50mM Tris pH 8.0, 150 mM NaCl, 1 mM EDTA, 1 mM DTT). Cleaved protein was collected in the eluate and the protease separated from the eluted proteins using a Glutathione Sepharose (Cytiva 17513201) column collecting in TNE buffer (50mM Tris pH 8.0, 150 mM NaCl, 1 mM EDTA). Proteins were dialyzed against suitable buffer, concentrated using centrifugal filter units (Millipore Sigma UFC9010), and depleted of endotoxin with Pierce™ High-Capacity Endotoxin Removal Spin Columns (Thermo Scientific 88274). Protein concentration was determined by BCA Protein Assay (Thermo Scientific 23227) and verified by Coomassie SDS-PAGE.

**Purification of recombinant soluble PGAM5 proteins**

PGAM5 plasmids were transfected in BL21 Rosetta competent *E. coli* cells for expression and plated on LB+Kanamycin^25ug/ml^ plate and left overnight at 37°C. Next day, 25ml of LB+Kanamycin^25ug/ml^ was inoculated with a few bacterial colonies from this overnight plate and left in a shaker incubator (37°C, 250rpm) overnight. Next day, 10ml of this culture was used to inoculate 500ml of LB+Kanamycin^25ug/ml^ (37°C, 250rpm) for a few hours until the OD reached 0.6-0.8. Protein expression was induced by adding 0.5mM IPTG and incubating for 18hrs (18°C, 225rpm). Cells were harvested by centrifugation and re-suspended in 15ml of lysis buffer (50mM Hepes-KOH pH7.5, 500mM NaCl, 5mM Imidazole, 5% glycerol with fresh 0.5mM TCEP, 1XPIM (200X: 976ml of H_2_O, 20ml of 1.5mg/ml-Aprotinin, 2ml of 10mg/ml-Pepstatin, 2ml of 10mg/ml-Leupeptin, 0.5mM PMSF). Cells were then sonicated (60% power, 3min/cycle, total 4-5 cycles) in the presence of 40mg/ml of DNAse. Supernatant was collected and incubated with 1ml of 50% Nickel(II)-NTA agarose (Ni^2+^-beads) (rock, 4°C, 2-3 hours). Ni^2+^-beads/supernatant mixture was settled in a column. The Ni^2+^-beads and bound His_6_-tagged PGAM5 proteins were washed with lysis buffer (10mM imidazole pH 7.4 with 0.5mMTCEP, 1XPIM, 0.2mMPMSF). Protein was eluted with lysis buffer containing higher imidazole concentration (250mM imidazole with 0.5mM TECP). Protein was concentrated to OD280 ~10-15 by using MWCO 10kDa concentrator and quantified using BCA assay. The purified proteins were dialyzed in 1X PBS, pH 7.4 and aliquots of the protein solution were frozen in liquid nitrogen and stored at -80°C after adding glycerol to a final concentration of 20% (v/v), for subsequent experiments.

**Isothermal Titration Calorimetry**

All ITC experiments were performed on a MicroCal PEAQ-ITC system at OU Protein Production and Characterization Core. For experiments involving APP constructs expressed in HEK293T cells, the purified PGAM5 and sAPPα constructs were dialyzed into a buffer containing 20 mM Na-HEPES pH 7.0, 150 mM NaCl, and 5 mM MgCl_2_. Dialyzed samples were concentrated and degassed prior to the experiment. In these experiments, sAPPα were placed in the sample cell, with matching buffer in the reference cell. PGAM5 constructs were loaded into the syringe and injected into the sample cell in a series of 1 μL injections at 25°C. The raw ITC data were fitted to a single binding site model using the MicroCal PEAQ-ITC analysis software provided by the manufacturer.

**Primary Mouse Astrocyte Culture**
Primary astrocytes were isolated from whole brains of postnatal day 0-5 (P0-P5) C57BL/6J mice using a differential adhesion-based protocol^58^. All dissections were performed under sterile conditions. Brains were rapidly removed, transferred to ice-cold Hank’s Balanced Salt Solution (HBSS), and meninges were carefully stripped to minimize contamination from fibroblasts and endothelial cells. Cortical tissue was mechanically dissociated and enzymatically digested to generate a single-cell suspension. Briefly, 0.05% DNase I was added to the tissue, followed by gentle trituration 10–15 times. Subsequently, 0.05% trypsin was added, and the tissue was further triturated 20–30 times using a 10 mL pipette before incubation at room temperature (22–25 °C) for 20 min. Cells were centrifuged for 10 min at 150×g, resuspended in complete Dulbecco’s Modified Eagle Medium (DMEM) supplemented with 10% fetal bovine serum and penicillin–streptomycin, and plated. The cells were cultured in the astrocyte media and maintained at 37°C with 5% CO₂. Media was changed on day 2 and subsequently every 2–3 days until they reached desired confluency. These were then either used for experiments or flash frozen for future use.

**qPCR**

Total RNA was extracted from WT and APP KO mice primary astrocytes using the Direct-zol micro-RNA isolation kit (Zymo Research) and quantified using a Nanodrop 1000 spectrophotometer. 0.25 μg RNA was used to synthesize cDNA using qScript (Quanta) according to manufacturer instructions in a BioRad T100 thermocycler. Real-time quantitative PCR was performed using PowerUp™ SYBR^®^ Green PCR Master Mix (Applied Biosystems) and the primers for the genes- *Hmox1, Nqo1, Gclc, Gclm*, *Txnrd1* and *Actb.* The mRNA levels of target genes were normalized to *ActB* by untreated WT sample values.

**Respiratory measurements with isolated mitochondria**

Freshly isolated mouse brain mitochondria were resuspended in mitochondrial resuspension buffer (250 mM mannitol, 5 mM HEPES, pH 7.4, and 0.5 mM EGTA) as previously described.^57^ The protein quantification was done via BCA assay. Mitochondrial respiration measurements were performed according to established protocols.^59^ Briefly, mitochondria were diluted to a final concentration of 0.25 mg/mL in respiration buffer containing 210 mM mannitol, 70 mM sucrose, 10 mM MOPS, and 5 mM KH₂PO₄ (pH 7.4). Substrate-driven respiration was assessed using either 0.1 mM pyruvate plus 1 mM malate, 10 mM glutamate plus 1 mM malate, or 1 mM succinate. Oxygen consumption was measured using a fluorescence lifetime–based dissolved oxygen monitoring system (Instech) at 20°C. State 3 respiration was initiated by the addition of ADP (0.5 mM) at 2 min, and oxygen consumption was monitored until complete conversion of ADP to ATP, corresponding to state 4 respiration.

**ETC enzyme activity assays**

NADH oxidase activity was measured as previously described.^60^ Briefly, isolated and protein quantified mitochondria were snap frozen in 25 mM MOPS (pH 7.4) at 0.025 mg/mL and kept in liquid nitrogen until ready to be used. To this frozen-thawed solution, 150 μM NADH was added and the rotenone-sensitive rate of NADH oxidation (ε_340_ = 6200 M^−1^ cm^−1^) in the presence of 10mM KCl was measured spectrophotometrically via an Agilent 8453 diode array UV−Vis spectrophotometer. To measure Complex 1 activity, 50nM antimycin A and 100uM UQ1 were added to the frozen-thawed mitochondria, followed by 150uM NADH. Complex I activity was then monitored spectrophotometrically as the rate of NADH oxidation. Rotenone (100 nM) was used as an inhibitor to ensure NADH utilization was dependent on complex I activity.
